# Reliability and correlation of fMRI, ECoG and EEG during natural stimulus processing

**DOI:** 10.1101/207456

**Authors:** Stefan Haufe, Paul DeGuzman, Simon Henin, Michael Arcaro, Christopher J. Honey, Uri Hasson, Lucas C. Parra

## Abstract

Human brain mapping relies heavily on fMRI, ECoG and EEG, which capture different physiological signals. Relationships between these signals have been established in the context of specific tasks or during resting state, often using spatially confined concurrent recordings in animals. But it is not certain whether these correlations generalize to other contexts relevant for human cognitive neuroscience. Here, we address the case of complex naturalistic stimuli and ask two basic questions. First, how reliable are the responses evoked by a naturalistic audio-visual stimulus in each of these imaging methods, and second, how similar are stimulus-related responses across methods? To this end, we investigated a wide range of brain regions and frequency bands. We presented the same movie clip twice to three different cohorts of subjects (N_EEG_ = 45, N_fMRI_ = 11, N_ECoG_ = 5) and assessed stimulus-driven correlations across viewings and between imaging methods, thereby ruling out task-irrelevant confounds. All three imaging methods had similar repeat-reliability across viewings when fMRI and EEG data were averaged across subjects, highlighting the potential to achieve large signal-to-noise ratio by leveraging large sample sizes. The fMRI signal correlated positively with high-frequency ECoG power across multiple task-related cortical structures but positively with low-frequency EEG and ECoG power. In contrast to previous studies, these correlations were as strong for low-frequency as for high frequency ECoG. We also observed links between fMRI and infra-slow EEG voltage fluctuations. These results extend previous findings to the case of natural stimulus processing.

## Introduction

The most frequently-used functional neuroimaging techniques are functional magnetic resonance imaging (fMRI) and electroencephalography (EEG). Both are complementary in that fMRI provides high spatial but low temporal resolution, while the opposite is true for EEG. Invasive methods such as electrocorticography (ECoG), on the other hand, combine high temporal resolution with relatively high spatial resolution; however, such procedures are only used in small cohorts of neurological patients. ECoG, EEG and fMRI also differ in their underlying neurophysiological origin, are susceptible to different noise sources (e.g., artifacts, pathological activity), and generally differ in terms of their inherent signal-to-noise ratio (SNR). From a practical perspective, it is, therefore, of interest to quantify and map out the amount of task-related information that each imaging modality contains in order to decide on the appropriate technique for a particular study.

At the same time, it is an ongoing endeavor to reveal the relationships between brain imaging techniques in order to better understand the physiological foundations underlying these methods. There have been significant efforts to relate the electrical neural signal measured with EEG/ECoG (Buzsáki et al., 2012) to the hemodynamic blood-oxygen-level dependent (BOLD) signal captured by fMRI (Logothetis, 2003). In respective experiments, imaging modalities are linked either through simultaneous recordings or through separate recordings tied together by a common task. Simultaneous fMRI–EEG recordings in human have been performed in Laufs et al. (2003); Moosmann et al. (2003); Ritter et al. (2009); Ritter and Villringer (2006); Scheeringa et al. (2008, 2011, 2016). Simultaneous fMRI–ECoG recordings are only available in animals (Logothetis et al., 2001; Magri et al., 2012; Niessing et al., 2005) with a recent exception in human (Carmichael et al., 2017). Most human studies have relied instead on a common task (Harvey et al., 2013; Hermes et al., 2012; Mukamel et al., 2005; Nir et al., 2007; Winawer et al., 2013). The main findings of these studies are that high-frequency power in the ECoG correlates positively with fMRI, while low frequencies correlates negatively with fMRI. These effects may, however, not be uniformly distributed across cortical brain structures, and can display frequency-dependent spatial variations (Harvey et al., 2013; Scheeringa et al., 2008, 2009). The observed correlations may also depend on the specific task (Maier et al., 2008; Muthukumaraswamy and Singh, 2009). Recent evidence suggests that the correlations between hemodynamic and electrical activity can have a non-neuronal physiological origin (Mateo et al., 2017), which can be decoupled from task-related neural processing (Winder et al., 2017). Thus, it is not certain whether the results of previous studies apply also in more realistic stimulus condition in humans, nor is it clear which of these relationships persist once controlling for physiological confounds unrelated to the brain functions under study.

The link between scalp EEG and invasive electrical recordings has been explored predominantly in non-human primates using simple visual stimuli. While visually evoked gamma activity generally correlates in the two imaging modalities (Whittingstall and Logothetis, 2009), there appears to be a complex relationship between lower frequency activity in the EEG and high-frequency intracranial activity (Snyder et al., 2015). In human, the link between ECoG and EEG has been explored mostly in the context of epileptic activity, but we are not aware of a systematic analysis of different frequency bands. Such an analysis may be warranted, given that results on visual evoked gamma activity, for instance, do not seem to readily extend to humans (Juergens et al., 1999) and likely depend on the stimulus (Scheeringa et al., 2016).

Here, we consider the case of a complex audio-visual movie stimulus and ask two basic questions that have not been sufficiently addressed in the previous literature. First, how reliable are the responses evoked by such a stimulus in EEG, fMRI and ECoG, considering the cohort sizes that are typically available in respective studies? And, second, to what extent are these three imaging modalities reflecting the same stimulus-related brain activity?

To study these questions, we analyzed correlations within and between EEG, fMRI and ECoG recordings across different brain structures and frequency bands. Data were acquired from three different cohorts of subjects within separate EEG, ECoG and fMRI studies, in which subjects watched an audiovisual movie. The richness of the movie stimulus thereby ensured that not only auditory, visual and multi-sensory systems were engaged in the viewing task, but also a host of higher-level cognitive functions such as language, memory, and attention. Using the same movie for all subjects allowed us to temporally align data of different recordings, and to quantify their similarity using correlations (Hasson et al., 2010, 2004). Relationships between imaging modalities were studied in a common anatomical space in terms of correlations between signals of different subject cohorts, here referred to as *inter-method correlations* (IMC). Additionally, the presence of two repeated viewings within each subject allowed us to assess the SNR of each imaging modality in terms of its repeat-reliability, which we quantified here in terms of interviewing correlations (IVC). The use of IMC/IVC thereby ruled out correlations induced by physiological or artifactual fluctuations not related to the task, which would be present in concurrent multi-modal recordings. To allow meaningful comparisons of correlations across methods, we harmonized the spatial and temporal scales of the different datasets using spatial co-registration and the standardization of correlation coefficients using data-dependent null distributions. Our study represents the first comprehensive quantification of the stimulus-related brain activity that is expressed within and shared between three important functional brain imaging modalities.

## Results

Three separate cohorts of subjects were presented with a 325 s long segment of a feature film (Dog Day Afternoon; previously used by Honey et al., 2012) two times, and neural activity was recorded with EEG (N = 45 subjects), fMRI (N = 11) and ECoG (N = 5). We analyzed correlations of ‘raw’ broad-band signals of all imaging methods as well as of EEG/ECoG power fluctuations in five frequency bands: θ (4–8 Hz), α (8–12 Hz), β (12–28 Hz), γ (28–56 Hz) and the high frequencies (HF, 64–116 Hz). To quantifying the effects that can be resolved from entire cohorts of realistic sizes, we performed grand-averaging (GA) of the EEG and fMRI data after spatial normalization. In ECoG, averaging across subjects is not possible, as electrode montages differ between subject; we, therefore, combined the ECoG channels of all subjects into a single dataset (without averaging). For comparison we also present fMRI and EEG results obtained on single subjects.

### Reliability of responses within imaging methods

To assess the *repeat-reliability* of stimulus-related brain activity for each method, we calculated the correlation of the continuous neural responses across the two renditions of the stimulus, which we refer to as the inter-viewing correlation (IVC). Correlations were assessed separately for each anatomical location using Pearson’s *r*, and their statistical significance was assessed by comparing observed values to a null distribution obtained from surrogate data. The resulting *z*-scores correct for unequal variances of the estimated correlation coefficients that arise from different sampling frequencies and data-dependent auto-correlation spectra. We interpret *z*-scores as objective measures of how much the observed correlations stand out against random fluctuations in the data. They can be directly compared across different imaging methods and differing sampling rates, whereas *r*-values are sensitive to the sampling rate and spectral content of the signals.

#### 2.1.1. Comparable repeated-reliability when averaging over typical cohort sizes

We observed strong IVC of neural activity for all three imaging methods (*r*-values in Fig. 1, and corresponding *z*-scores to determine significance in Fig. 2). The grand-average fMRI BOLD signal (GA-fMRI) reached a maximum IVC of r_max_ = 0.79, where strongest correlations were observed in temporal, parietal, occipital and fronto-temporal areas known to implement auditory and visual processing hierarchies (Fig. 1 A).

**Fig. 1.**
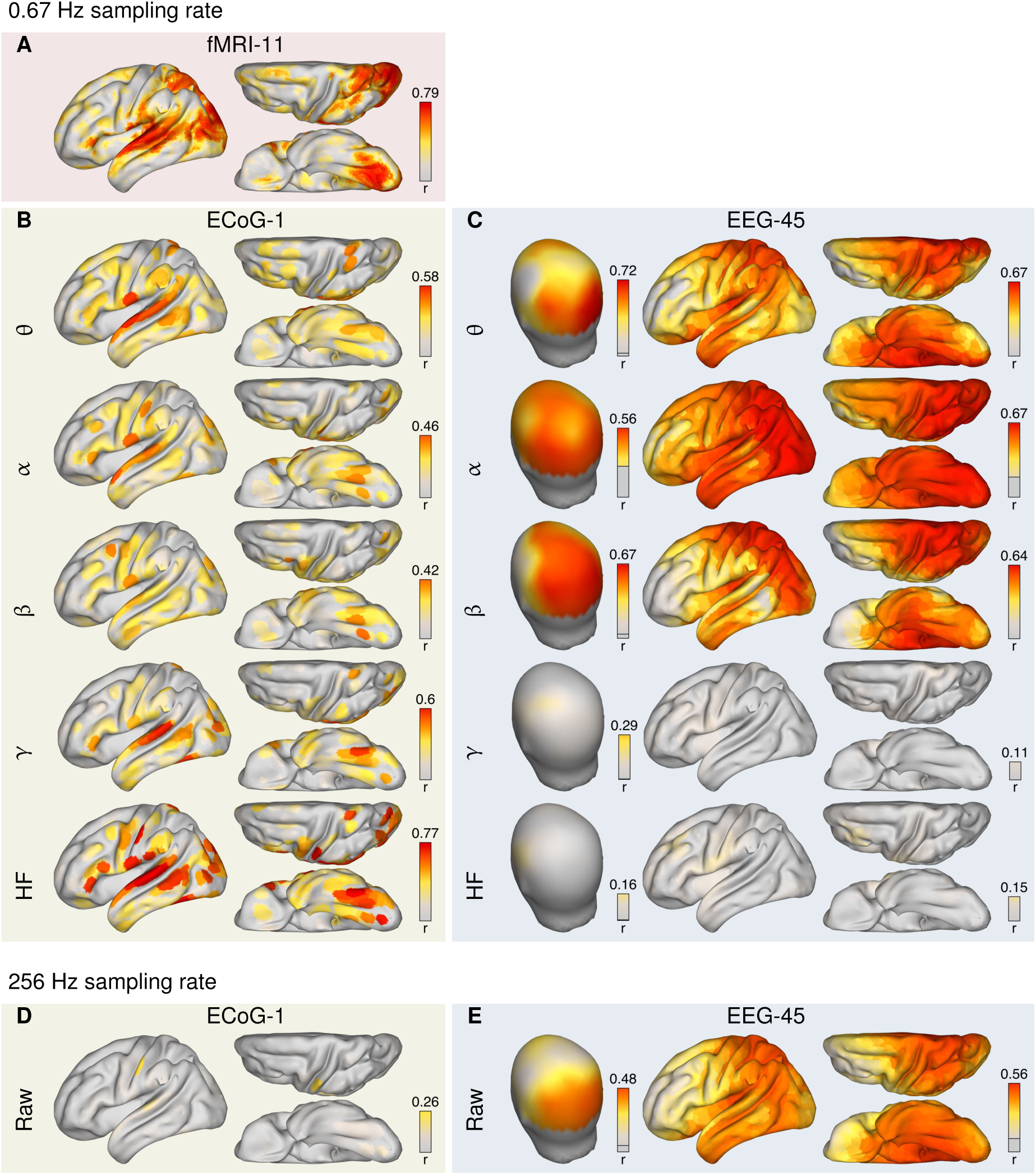
Repeat-Reliability of stimulus-related brain activity measured with EEG, ECoG and fMRI BOLD. measured as inter-viewing correlations (IVC), *r* of signals acquired during two separate viewings of a 325 s long audio-visual movie stimulus. EEG data were analyzed at 32 sensors and 2,004 cortical locations after source reconstruction by eLORETA. fMRI data were analyzed at 16,037 cortical locations, while ECoG data were analyzed at 511 cortical locations representing the pooled set of electrodes in five subjects. ‘Raw’ broad-band EEG and ECoG voltage fluctuations were analyzed at 256 Hz sampling frequency (panels **D, E**). In addition, the instantaneous log-power of EEG and ECoG oscillations in the following frequency bands were analyzed at the same sampling frequency as fMRI (0.67 Hz, panels **A–C**): θ (4–8 Hz), α (8–12 Hz), β (12–28 Hz), γ (28–56 Hz) and high-frequency (HF, 64–116 Hz). EEG data of 45 subjects and fMRI data of 11 subjects were grand-averaged before IVC analysis. IVC was measured in terms of Pearson correlation, *r*, and plotted onto the outer head surface (sensor-space EEG) or the smoothed cortical surface of the left brain hemisphere (ECoG, fMRI, source-space EEG, shown from left, top, and bottom). Numbers above the color bars indicate maximal correlation values. As ECoG data do not provide a full coverage of the cortex (see Fig. S1), gray areas mark the absence of any electrode within 12 mm distance.

**Fig. 2.**
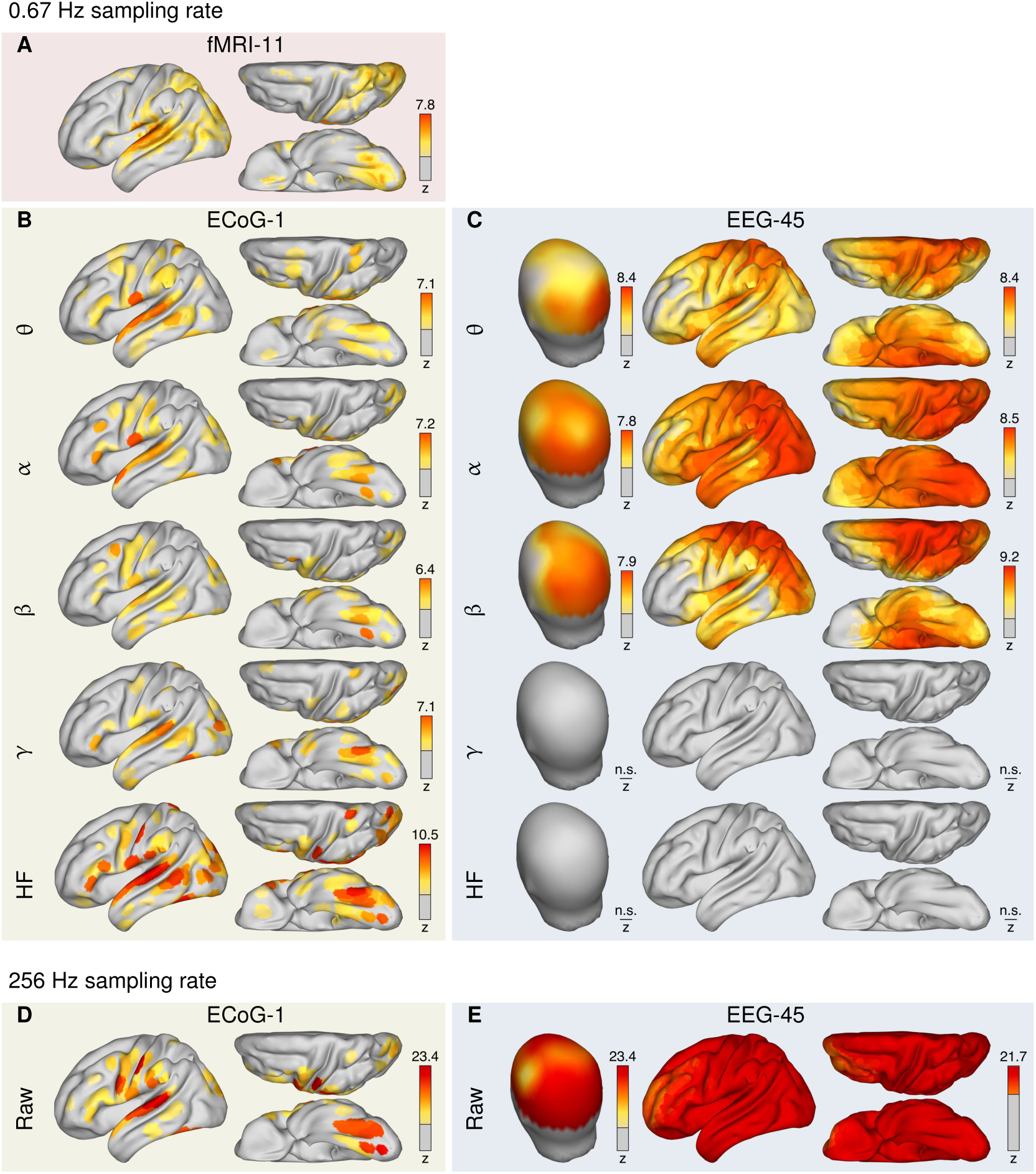
Statistical significance of inter-viewing correlation (IVC). Observed correlations, *r*, were standardized based on the mean and standard deviation of a null distribution established from phase-randomized surrogate data, yielding a standard normal distributed *z*-score for each scalp/cortical location. *z*-scores are shown in color if they indicate statistical significance at level *q* < 0.05 (one-tailed test, FDR corrected). The figure shows that even small IVC values observed for ‘raw’ broad-band EEG/ECoG voltage fluctuations (c.f., panels D, E to corresponding panels of Fig. 1) are highly significant when compared to surrogate time series with identical autocorrelation spectrum but no IVC. Numbers above the color bars indicate maximal correlation values.

Strong IVCs of single-subject ECoG band-power traces were observed in very similar areas in all studied frequency bands (Fig. 1 B). In contrast to fMRI, significant IVC were present in central sensori-motor areas, while the lack of ECoG electrode coverage in parietal regions (see Fig. S1) prevented IVC analysis in these regions. The strongest IVC were in the same range as those observed for GA-fMRI (r_max_ = 0.77), and were observed in the superior temporal gyrus for high-frequency oscillations.

GA-EEG band-power in the lower frequency bands (θ, α, and β) reached maximal IVC levels only slightly below what is observed for ECoG and GA-fMRI (r_max_ = 0.72, see Fig. 1 C). The levels of IVC were comparable when computed on scalp sensor data and on cortical sources estimated using a linear solution to the EEG inverse problem (Pascual-Marqui, 2007, eLORETA). In both cases, the topography of IVC showed a broad global pattern, with low IVC values observed only in fronto-temporal regions. High-frequency and γ-band activity has negligible IVC, speaking to the susceptibility of the EEG signal to movement artifacts and the relative weakness of coherent brain activity reaching scalp sensors in these frequency ranges.

For the raw broad-band EEG and ECoG voltage fluctuations (Fig. 1 C, D), IVC reached maximal values of *r*_max_ = 0.56 in GA-EEG and *r*_max_ = 0.26 in ECoG. The topography of raw EEG was similar to the topography obtained with EEG θ-band power, while the raw-ECoG topography closely resembled the ICV distribution observed in HF-ECoG.

To demonstrate the effects of grand-averaging, we also computed IVC on single subjects (Fig. S2). Without averaging, the maximum IVC in fMRI reached only r_max_ = 0.39 with a similar spatial distribution (panel A). Grand-averaging was particularly effective for EEG (compare Fig. 1 C, E with Fig. S2 B, C). Without grand-averaging, IVC for single-subject EEG only reached values around 0.1 (notice that the color map has been enhanced to show these smaller value). These smaller IVC are consistent with previous reports in EEG and MEG (Dmochowski et al., 2012; Lankinen et al., 2014).

#### 2.1.2. Repeat-reliability is highly significant in all modalities, independent of sampling rate

To ensure compatibility with fMRI and previously published ECoG results (Honey et al., 2012), the IVCs computed for EEG and ECoG band-power traces shown in Fig. 1 B, C were obtained on data reduced to the fMRI sampling rate of 0.67 Hz. Note, however, that correlation values typically depend on the spectral content of the signal and should not be directly compared across different sampling rates. Similarly, maximum values should not be directly compared between differing sampling rates as they depend on the number of samples. Indeed, when evaluated at a higher sampling rate of 256 Hz, IVCs of the band-passed powers dropped to a maximum of *r*_max_ = 0.23 for ECoG and *r*_max_ = 0.53 for EEG (compare Fig. S3 A, B with Fig. 1 B, C).

The *z*-scores we calculated (Fig. 2) measure how much the observed IVCs depart from values obtained with random signals that have the same spectral content as the original signals (see Statistical significance of correlations). Significant z-scores were obtained for all three imaging modalities across large portions of the cortex (Figs. 2). Surprisingly, by far the strongest values were observed for the raw GA-EEG and single-subject ECoG (z_max_ = 23.4 for both, see Fig. 2 D, E) despite the moderate to weak *r*-values (compare with Fig. 1 D, E).

The dependence of the *r*-values on sampling rate noted above largely disappear for the *z*-scores (compare Figs. 2 B, C and S4 A, B). An explanation for this stability is that the noise (non-stimulus-related activity) above 0.33 Hz diminishes inter-viewing correlations as well as correlations that arise under the null hypothesis, limiting the variance of the latter. Removing that noise through low-pass filtering increases IVC, but to the same extent also correlations that could arise under the null hypothesis, leading to the same level of significance (see also Fig. S5). If anything, maximal *z*-scores were higher for EEG power fluctuations in the θ-, α-, and β-bands when sampled at 256 Hz (*z*_max_ = 15.9) as compared to 0.67 Hz (*z*_max_ = 9.2). The same was observed for HF-ECoG oscillations (*z*_max_ = 15.3 for 256 Hz compared to *z*_max_ = 10.5 for 0.67 Hz sampling rate). This increase in correlation suggests that stimulus-related amplitude fluctuations in these data are predominantly occurring at frequencies below 0.33 Hz (see Fig. S5). However, down-sampling reduced *z*-scores because a substantial amount of reliable activity occurs above 0.33 Hz, as verified by applying a spectrally-resolved inter-viewing coherence analysis (Fig. S5).

### Similarity of responses between imaging methods

To assess the *similarity* of responses between imaging methods, we measured the *inter-method correlation (IMC)* after spatial co-registration (see Spatial registration and Mappings between fMRI, ECoG and EEG spaces for technical details). As before, results for fMRI and EEG were computed with the signal averaged across subjects (grand-average). We observed significant correlations of stimulus-related responses between all three neuroimaging methods (IMC are shown in Fig. 3 and the corresponding *z*-scores in Fig. 4). IMC were strongest between the GA-fMRI and ECoG band-power with absolute values exceeding *r*_max_ = 0.61 in all frequency bands (Fig. 3 A). ECoG power in the lower frequency bands (θ, α, β) was negatively correlated with BOLD in virtually all studied areas with large IMC (θ: *r*_min_ = −0.72, α: *r*_min_ = −0.72, β: *r*_min_ = −0.70). In contrast, γ- and HF-ECoG power in the same areas was positively correlated with fMRI (γ: *r*_max_ = 0.61), HF: *r*_max_ = 0.72). Negative correlations between ECoG γ-power with GA-fMRI were not significant (Fig. 4 A).

**Fig. 3.**
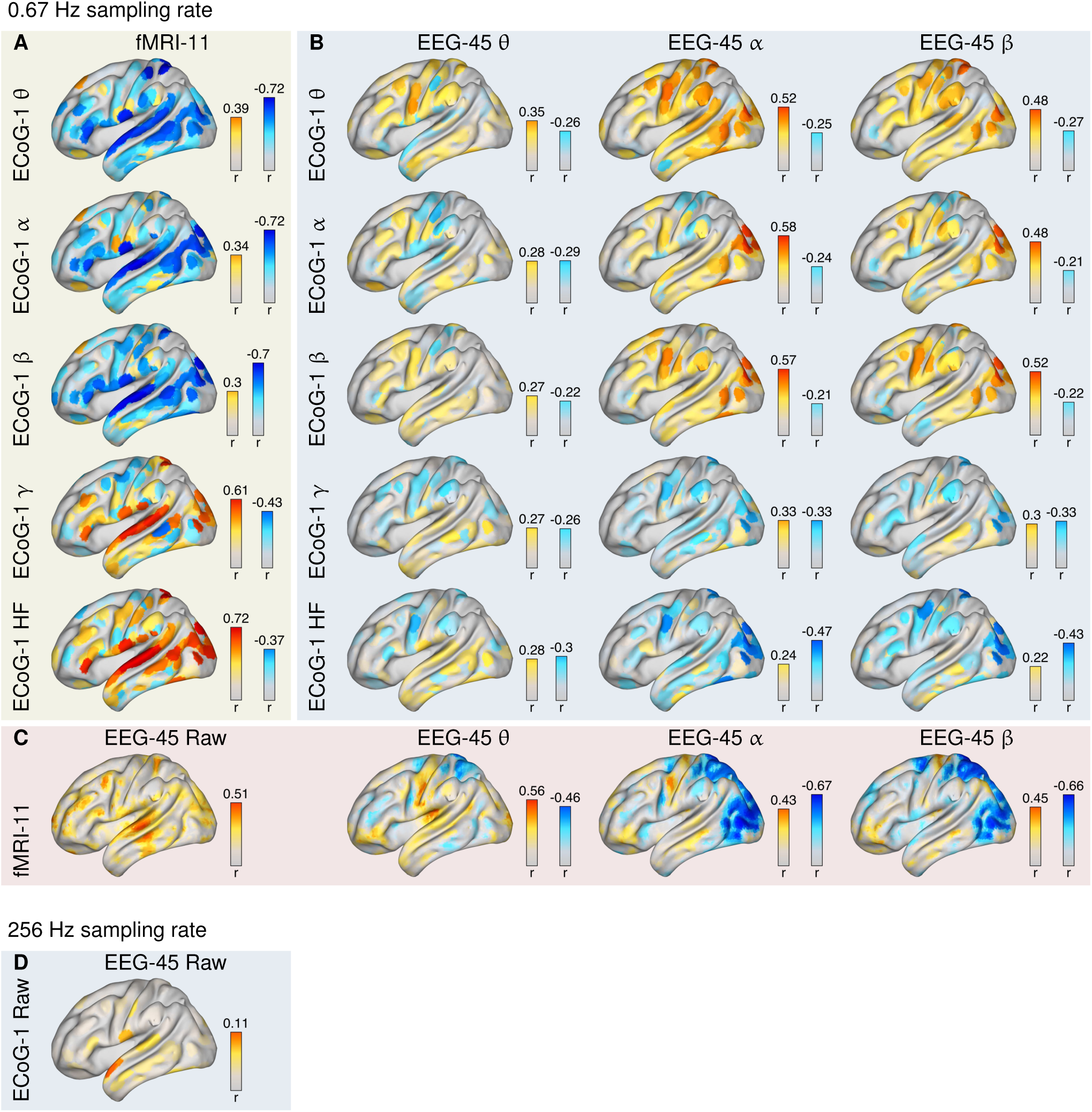
Similarity of stimulus-related brain activity between EEG, ECoG and fMRI BOLD as measured in terms of inter-method correlations (IMC) between co-localized single-subject ECoG, grand-average EEG and grand-average fMRI BOLD activity. IMC was assessed in terms of Pearson correlation, *r*, between ‘raw’ broad-band EEG and ECoG voltage fluctuations at 256 Hz sampling rate (panel **D**, notice the different color scale compared to panels A–C as well as between fMRI and raw EEG, between fMRI and EEG band-power, between between fMRI and ECoG band-power, and between EEG and ECoG band-power (panels **A–C**) at 0.67 Hz sampling rate. Frequency bands are the same as in Fig. 1. Matching of co-localized ECoG electrodes and cortical fMRI voxels was based on Euclidean distance. Source-reconstructed EEG activity (eLORETA) was computed for each fMRI voxel location. Numbers above color bars indicate maximal correlation values.

**Fig. 4.**
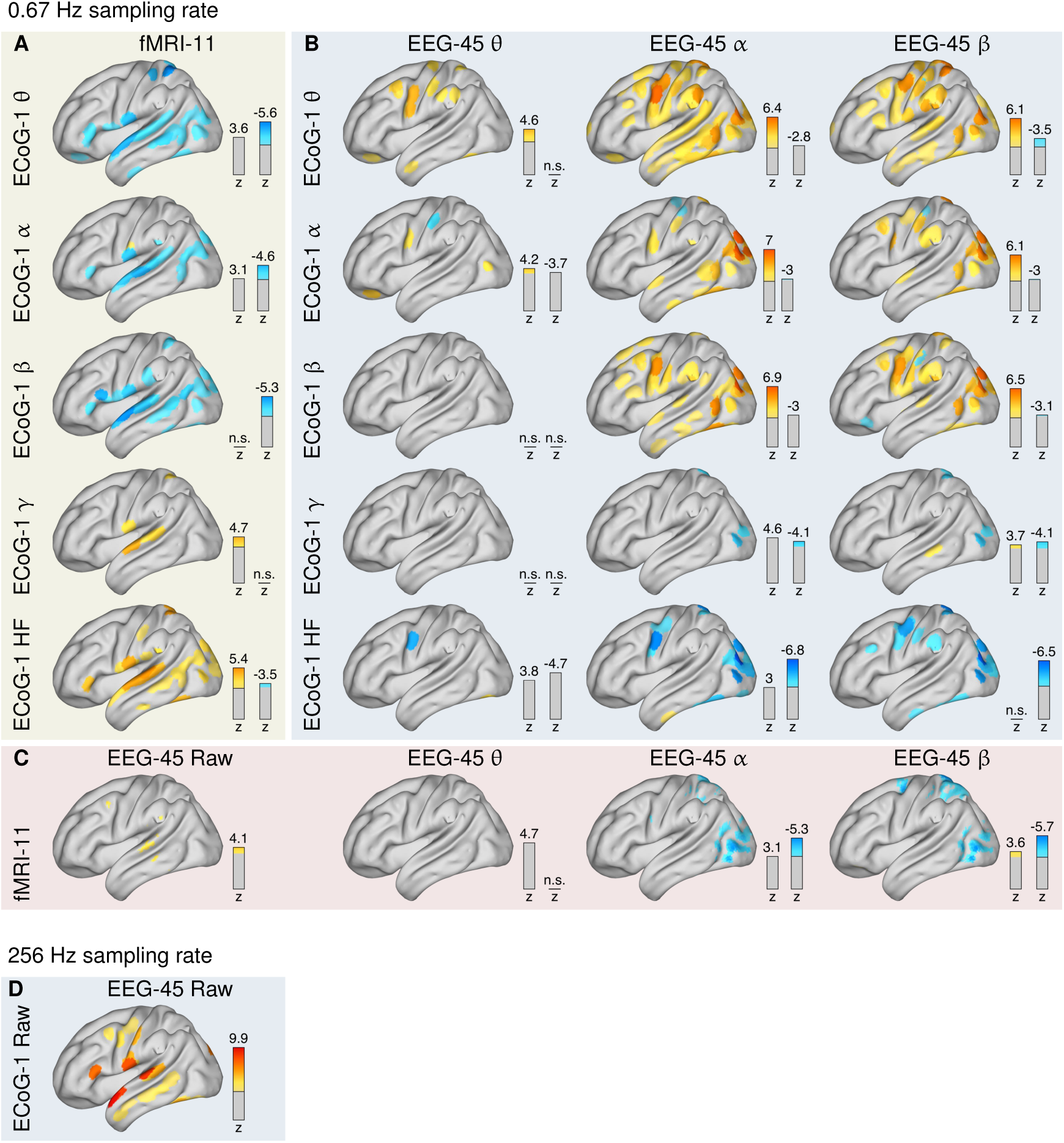
Statistical significance of inter-method correlations between single-subject ECoG, grand-average EEG and grand-average fMRI BOLD activity. Using a null distribution established from phase-randomized surrogate data, observed correlations were transformed into standard normal distributed z-scores. Significant z-scores (using FDR control at level *q* = 0.05) are drawn in color.

EEG low-frequency band-power was positively correlated with ECoG low-frequency power but negatively correlated with high-frequency ECoG (Fig. 3 B). Positive correlations in the low-frequency bands (θ, α, β) extended over central and temporal cortices, and are strongest in occipital cortex (in the range of *r*_max_ = 0.48 to 0.58. For γ and HF ECoG, correlations with low-frequency EEG were predominantly negative and again strongest in occipital cortex (*r*_max_ = 0.47).

EEG band-power in the lower frequencies shows diverse spatial effects relative to fMRI (Fig. 3 C). EEG α- and β-power fluctuations correlated negatively with fMRI in occipital and parietal areas (*r*_min_ = −0.67 for both). In addition, a negative correlation in parietal areas was found for θ-power (*r*_min_ = −0.46). Yet, EEG θ-and α-power correlated positively with fMRI BOLD in the temporal (θ: *r*_max_ = 0.56) and sensory-motor areas (α: *r*_max_ = 0.43); with weak corresponding *z*-scores Fig. 4 C). A possible explanation for this mixed finding is the fact that fMRI itself exhibits anti-correlated activity for this stimulus (Fig. S6): occipital fMRI is weakly negatively correlated with fMRI in temporal cortices and a portion of sensory-motor cortex (*r*_max_ = −0.18). When analyzing ‘raw’ EEG and ECoG evoked responses (not powers but phase sensitive signals), we observed significant correlations between the fMRI signal and raw infra-slow EEG below 0.33 Hz (Fig. 3 C), predominantly in temporal areas (*r*_max_ = 0.51). There was also highly significant correlation of the raw ECoG with raw EEG fluctuations over temporal areas (Fig. 3 D, *r*_max_ = 0.11). A similar IMC analysis between raw ECoG and fMRI was not possible due to a high-pass filter of 0.6 Hz that was applied to the ECoG data at recording time.

Finally, note that correlations between fMRI, EEG band-power and ECoG band-power were assessed above on data sampled at 0.67 Hz so that correlations and in particular maximum values can be compared between methods (Fig. 3). When repeating these analyses at 256 Hz sampling frequency, we obtain similar results (Figs. S7–S8).

## Discussion

In this work, we addressed two basic questions concerning fMRI, ECoG and EEG. First, how reliably can each imaging modality measure responses elicited by naturalistic stimuli? Second, how similar are these stimulus-responses in the three modalities? Our analyses identified brain areas related to the processing of a complex audio-visual narrative in temporal, occipital, parietal and sensori-motor areas, in line with previous reports (Honey et al., 2012; Jacques et al., 2016). Reliable stimulus-related activity in these areas was found not only in fMRI BOLD signals and high-frequency power of the ECoG, but also in low-frequency EEG and ECoG rhythms as well as in raw broad-band EEG and ECoG voltage fluctuations. The regions involved in stimulus-related processing are strikingly consistent between fMRI and ECoG, suggesting that both methods pick up similar brain responses, as previously established. This finding was further corroborated by studying correlations between methods. In line with results obtained in simultaneous fMRI-ECoG recordings in animals (Logothetis et al., 2001; Magri et al., 2012; Niessing et al., 2005) as well as in human (Carmichael et al., 2017), we observed positive correlations between fMRI and high-frequency ECoG activity and negative correlations with low-frequency ECoG. We also observed negative correlations between EEG α band-power and BOLD in parietal, occipital and rolandic areas in line with previous reports on simultaneous recordings (Laufs et al., 2003; Moosmann et al., 2003; Ritter et al., 2009; Scheeringa et al., 2009). Overall, these results are in line with the accepted notion that fMRI and HF ECoG capture similar activity, namely, neuronal firing (Logothetis, 2003; Manning et al., 2009; Nir et al., 2007), yet low-frequency oscillations are qualitatively different. α-band activity in particular is often said to reflect top-down inhibitory processes (Halgren et al., 2017; Klimesch et al., 2007) that suppress neuronal firing, and is thus negatively correlated with HF activity (Spaak et al., 2012). Importantly, and contrary to previous work (Magri et al., 2012), here we find that these lower frequencies are just as much correlated with fMRI BOLD as high frequencies for the studied natural-stimulus viewing task.

Interestingly, we found both positive and negative correlations between fMRI and EEG band power within the θ and α-bands of the EEG. Negative correlations were clearly observed in occipital cortex in the α, while correlations were positive in temporal and motor cortices in the β-band. Further analyses showed that this mixed sign could be the result of anti-correlated fMRI activity in visual and auditory cortex, which may reflect a competition of cognitive processes for attentional resources during processing of the multi-modal stimulus (Shomstein and Yantis, 2004).

A noteworthy result of the present study is that we identified strong stimulus-induced correlations of the BOLD signal and the infra-slow EEG signals below 0.33 Hz (Fig. 3, panel B). Previous links between hemodynamic responses and infra-low EEG had been associated with resting-state fluctuations (Hiltunen et al., 2014), the slow-cortical potential (He and Raichle, 2009), and arterial blood pressure (Nikulin et al., 2014). Most studies in this field used simultaneous recordings; correlated activity could therefore be the result of common physiological or artifactual confounds such as heart beat, breathing, etc. Here, we could exclude the presence of any confound unrelated to the cognitive task. The observed correlations between infra-slow EEG and fMRI were most pronounced over temporal areas, suggesting auditory and speech perception processes as drivers, though this may be specific to the present audiovisual video stimulus.

### Benefits and limitations of inter-subject correlations

Our analyses relied on correlations between separately-acquired but synchronized recordings within and across imaging methods. While this framework is limiting in that it requires identical timings of all relevant events, it does allow us to assess the similarity of signals from different imaging methods without requiring multi-modal recordings. An advantage of this approach is that one can prevent the technical complications and significant signal-loss resulting from simultaneous recordings. More importantly, the approach prevents task-irrelevant physiological processes from driving the results (Hasson et al., 2004; Mukamel et al., 2005), a concern that has been recently highlighted for fMRI, in particular during ‘resting state’ (Winder et al., 2017). The study of cognitive processes using inter-subject correlations is well established in fMRI (Chen et al., 2017; Jääskeläinen et al., 2008; Mantini et al., 2012), ECoG, (Honey et al., 2012), EEG (Cohen and Parra, 2016; Dmochowski et al., 2012; Ki et al., 2016), and MEG (magnetoencephalography, Lankinen et al., 2014). As with those studies, all correlations reported here exclusively relate to brain processes involved in processing the audio-visual narrative. The spatial distribution of reliable brain activity reported here must, therefore, be interpreted exclusively in terms of that cognitive task, and should not be mistaken as representative of all brain signals captured by the different imaging methods.

### Cohort size and impact of grand-averaging

EEG and fMRI data presented here were averaged across subjects before assessing the reliability of the task-related responses they contain. This form of ‘grand-averaging’ is well established to analyze evoked potentials in EEG and has recently also been used in fMRI (Schmaelzle et al., 2017). We did not average across subjects in ECoG due to inconsistent electrode montages in each of the five subjects. Using grand-averaging in EEG and fMRI, the SNR (or, repeat-reliability) could be increased to reach similar levels of reliability as in single-subject ECoG. Without grand-averaging, the observed reliability of fMRI was lower than in ECoG. This drop was even more pronounced in EEG. Note that the sample sizes used here are typical in research studies, and are indicative of the ease of obtaining these data for the different methods. The obvious disadvantage of averaging is the potential loss of spatial (and functional) resolution due to anatomical or functional misalignment across subjects, as well as the loss of subject-level information. Whether or not grand-averaging is a useful approach in a given neuroimaging study will depend on this trade-off between (functional) resolutions and signal-to-noise ratio as well as the general analysis goal.

Among the various signals analyzed here, raw EEG/ECoG times series (above 0.6 Hz), typically referred to as ‘evoked responses’, were somewhat less reliable across stimulus repetitions, but were highly significant compared to chance fluctuations (for the EEG, after grand-averaging). Raw stimulus-evoked EEG activity has traditionally been studied by averaging brain responses across many repetitions of a simple stimulus within highly controlled experiments (Luck, 2014). More recently it has been used with naturalistic stimuli by correlating activity between subjects (Cohen and Parra, 2016; Dmochowski et al., 2014; Ki et al., 2016). Recent evidence places the origin of these evoked responses to superficial cortical layers with broad spatial coherence (Halgren et al., 2017), suggesting a likely reason for why they dominate the scalp EEG. Here we found that the potential fluctuations evoked by the video stimulus correlate between EEG and ECoG, most strongly over temporal cortex. Given the large z-scores observed here, these signals may have been generally under-appreciated, as they remain largely unexplored in ECoG and generally in the context of continuous naturalistic stimuli.

### Identifying the most reliable time-scale

We analyzed amplitude fluctuations in EEG and ECoG at the original sampling rate of 256 Hz as well as at 0.67 Hz. Low-pass filtering to the lower sampling rate substantially increased inter-viewing and inter-method correlations. Slow fluctuations below 0.33 Hz were found to capture most of the stimulus-related signal in these amplitude traces. Interestingly, z-scores were found to be larger before low-pass filtering, while corresponding inter-viewing correlations increased. This suggests that z-scores are less sensitive to the power-spectrum of the stimulus-related and noise portions of the signal, and a drop in z-scores indicates that reliable activity has been removed.

### Comparing reliability between imaging methods

An important advantage of z-scores over r-values is that they allow direct comparison between data sets. For homogeneous datasets consisting of a fixed number of independent samples, there is a monotonous relationship between r-values and z-scores. Thus, comparisons of datasets in either of the two metrics lead to the same conclusion. This is not anymore the case if the number of samples differs between datasets, as correlations estimated on fewer samples are more variable and therefore reach statistical significance less easily. The same effect is caused by dependencies between samples, which lead to increased variance by decreasing the *effective* number of samples. Such dependencies are naturally present in all neurophysiological time series, and are expressed in dataset-specific autocorrelation spectra. Appendix A provides a numerical demonstration that z-scores as computed here achieve correct false-positive rates (p-values) despite different number of samples and autocorrelation spectra. Thus, z-scores are suitable to compare IVC in EEG, fMRI and ECoG despite differing numbers of samples and differing spectra of these signals.

In practice, the choice of the ‘right’ metric depends on one’s analysis goals. Strong correlations (if significant) are useful if one wants to predict one time course from another, where stronger correlations imply better predictability. This could, for example, be of practical relevance if one wanted to approximate ECoG activity from non-invasive EEG measurements. In contrast, if one is interested in statistical significance, then the relevant metric is the *z*-score. For example, one may want to test if IVC differs between task conditions. In those instances the relevant question is not how large or small the correlations are, but rather, how stable they are in the face of chance fluctuations. There, a high *z*-score is more relevant than a large *r*-value.

### Spatial heterogeneity

The EEG, ECoG and fMRI datasets studied here strongly differ in the number of simultaneously-acquired measurements, as well as in the spatial domains sampled by these measurements. EEG was measured at 32 scalp sites, whereas ECoG and fMRI were assessed at about 100 intra-cranial electrodes and thousands of brain voxels, respectively. In this light, one of the most interesting findings of this study is that EEG signals reached absolute levels of reliability that are comparable to levels achieved by ECoG and fMRI. A likely explanation for the competitive performance of EEG is again the larger number of subjects entering the grand-average, as discussed above.

Computation of inter-method correlations was hampered by different spatial coverages. fMRI signals were available from the entire brain, but only cortical activity was used for inter-method comparisons. EEG sensors pick up mixtures of activities of sources located all across the brain, and therefore often suffer from low SNR and the lack of straightforward anatomical localization. To increase SNR, and to visualize and relate EEG to ECoG and fMRI activity in a common anatomical space, scalp EEG data were mapped to the cortical surface using an inverse source reconstruction technique. The blurring observed in EEG source estimates could in principle be reduced by making prior assumption on the spatial focality of the underlying brain sources (e.g., Haufe et al., 2008, 2011). However, it was accepted here as being reflective of the low spatial resolution of the scalp-level data. ECoG electrodes were available at superficial cortical locations of the left hemisphere, with little coverage for large parts of the right hemisphere, the medial surface of the left hemisphere, and several larger patches of the occipital, parietal, central and frontal cortex of the left hemisphere. Inter-method correlations involving ECoG were thus only assessed at those locations.

### Future work

Two restrictions of the present study are that we assessed only linear relationships and only looked at co-localized anatomical structures. Future studies may assess task-related interactions between different brain structures (Simony et al., 2016) across imaging methods and use non-linear measures of functional connectivity in order to better characterize the mechanisms linking the neurophysiological phenomena picked up by different methods. Using partial correlation analysis and information theory, future studies may also attempt to dissociate task-related signals that are method-specific from signals that are reflected by multiple imaging methods. Specific multivariate techniques (e.g., Bießmann et al., 2010; Dähne et al., 2015; Dähne et al., 2014; Dmochowski et al., 2012; Lankinen et al., 2014; McIntosh and Lobaugh, 2004) could, moreover, be used to identify brain networks characterized by maximal IVC/IMC in optimal data-driven ways.

## Conclusion

Our results provide a comprehensive spatio-spectral account of the neural correlates of natural audio-visual stimulus processing in fMRI, ECoG and EEG, three of the most widely used neuroimaging methods in humans. All three methods reached similar levels of reliability when data were averaged across the subjects of each cohort. Correlations between methods confirmed prior findings of an opposing sign of high and low frequency electrical activity, perhaps indexing different neural mechanisms (direct neuronal firing in high-frequency ECoG and fMRI BOLD, inhibitory drive in low-frequency EEG/ECoG).

## Experimental procedures

ECoG data used in this study have been described in detail in Honey et al. (2012); we here largely recapitulate these details. The EEG data with 32 electrodes (*N* = 30 subjects) has previously been used by Dmochowski et al. (2017). The additional EEG dataset with 128 electrodes (*N* = 15 subjects) and the fMRI data have not been previously published.

### Study subjects

#### fMRI

Eleven subjects (six female; 20-35 years old) participated in the fMRI experiment. All subjects were in good health without history of psychiatric or neurological disorders and gave their informed consent to participate in the study and consent to publish in accordance with procedures approved by the Princeton University Institutional Review Board and with ethical standards set out by the Federal Policy for the Protection of Human Subjects (or ‘Common Rule’, U.S. Department of Health and Human Services Title 45 CFR 46). Subjects had normal or corrected-to-normal visual acuity. All subjects were experienced MRI subjects that were well trained to lay still during scans. There were no specific recruitment goals with respect to gender or race/ethnicity; however, fluency in English was required.

#### ECoG

The following description is quoted from (Honey et al., 2012): “Five patients (four female; 20–47 years old) experiencing pharmacologically refractory complex partial seizures were recruited via the Comprehensive Epilepsy Center of the New York University School of Medicine. Patients had elected to undergo intracranial monitoring for clinical purposes and provided informed consent both pre-and post-electrode implantation in accordance with National Institutes of Health guidelines administered by the local Institutional Review Board. For each patient, electrode placement was determined by clinicians based on clinical criteria. We focus here on patients with entirely or predominantly left-lateralized coverage, all of whom had left-lateralized language function.” We are excluding right-hemisphere electrodes available from two patients with bilateral coverage.

#### EEG

Data were obtained from two separate batches. In the first batch, thirty healthy subjects were recruited (fifteen male; 19–31 years old). Procedures were approved by the Western Institutional Review Board (Puyallup, WA). In the second batch, 15 healthy subjects were recruited from the campus of City College (nine male; 18–28 years old). All procedures for this cohort were approved by the Institutional Review Board of the City University of New York. Prior to the start of the experiments, all subjects gave written informed consent. There were no specific recruitment goals with respect to gender or race/ethnicity; however, fluency in English was required.

### Experimental setting

The audiovisual stimulus was a 325 s long movie clip selected from the 1975 commercial film Dog Day Afternoon (DDA, Lumet, 1975).

#### fMRI

subjects viewed two repetitions of DDA clips. In addition to original (intact) clips, two manipulated variants, in which the scene order was randomized either on a coarse or fine level, were presented. The order of presentation was fixed: intact, coarse, intact, fine, coarse, fine.

Presentation order was randomized across individuals. The Psychophysics Toolbox in MATLAB was used to display the movie clips and synchronize the movie onset with the MRI data acquisition. Audio for the movie was delivered via in-ear headphones. Movie clips subtended 20° horizontally and 16° vertically.

#### ECoG

subjects viewed DDA in alternation with two more movie clips (two presentations per clip) at bedside on a Mac-Book laptop located 40–60 cm from their eyes. PsychToolbox Extensions (Kleiner et al., 2007) for MATLAB (MathWorks, Natick, MA) were used to display the movies and trigger their onsets. Subjects viewed intact, coarse, and fine renditions of the clips. Presentation of each clip was preceded by a 30 s period in which subjects fixated on a central white square (< 1° visual angle) on a black background.

#### EEG

subjects of the first batch watched two repetitions of the intact movie interspersed with single presentations of the manipulated version: intact, coarse, fine, intact as used in Honey et al. (2012). In-house software (Neuromatters LLC, NY) was used for video playback. Only data recorded during presentation of intact stimuli (two viewings per subject and imaging method) were analyzed in the present study. subjects of the second batch watched the intact DDA clip twice, with each exposure separated by approximately 45 min, at the beginning and the end of an unrelated experiment. The video clip was presented via a custom version of mplayer (Mplayer media player, http://mplayerhq.hu), modified to generate parallel port triggers once per second.

### Signal acquisition

#### fMRI

Data were acquired with a 3T Skyra magnetic resonance imaging (MRI) scanner (Siemens, Munich, Germany) using a 16-channel head coil. All functional acquisitions used a gradient echo, echo planar sequence with a 64 square matrix (slice thickness of 4 mm, interleaved acquisition) leading to an in-plane resolution of 3 × 3 mm^2^ (field of view [FOV], 192 × 192 mm^2^, GRAPPA iPAT = 2, 27 slices per volume; repetition time [TR] = 1.5 s; echo time [TE] = 30 ms; flip angle = 72 degrees). High resolution structural scans were acquired in each scan session for registration to surface anatomical images (MPRAGE sequence; 256 × 256 matrix; 240 × 240 mm^2^ FOV; TR = 1.9 s; TE = 2.1 ms; flip angle = 9 degrees; 0.9375 × 0.9375 × 0.9375 mm^3^ resolution).

#### ECoG

Signals were recorded from 922 electrodes across all five subjects. Subdural arrays of platinum electrodes embedded in silastic sheeting (8 × 8 square grids, 4 × 8 rectangular grids, or 1 × 8 strips) were placed purely according to clinical criteria. Electrodes had an exposed diameter of 2.3 mm and were spaced 10 mm center-to-center. Depth recordings were not analyzed in the present study. Screws in the skull served as reference and ground. Signals were sampled at 30 kHz using a custom-built digital acquisition system (based on the open-source NSpike framework (L.M. Frank and J. MacArthur, Harvard University Instrument Design Laboratory, Cambridge, MA) that included a 0.6 Hz high-pass filter in hardware.

#### EEG

EEG data were recorded with a BioSemi Active Two system (BioSemi, Amsterdam, Netherlands) at a sampling frequency of 2,048 Hz. subjects of the first batch were fitted with a standard, 32-electrode cap following the international 10/20 system, while subjects in the second batch were fitted with a 128-electrode cap according to an equiradial system (Biosemi). Six additional electrooculogram (EOG) electrodes were placed around the eyes to record and allow for the removal of eye-movement artifacts. Sony MDR 7506 headphones were used for audio playback during video viewing for the first batch of subjects, whereas the second batch of subjects were presented the audio over a pair of studio monitors, each situated at a 45° angle to the subject (e.g., left/right side) and at a distance of approximately 1 m (Fostex, PM0.3 Active monitors, Tokyo, Japan).

For this investigation, we pooled the data of the two EEG batches. To this end, 32 out of 128 electrodes used in the second batch of recordings were matched with the 32 electrodes used in the first batch of recordings based on nearest Euclidean distance of their standardized locations (median distance: 6 mm, maximum: 12 mm). This yielded N = 45 recordings with 32 EEG and six EOG electrodes available in each.

### Spatial registration

All data were processed using in-house MATLAB code unless otherwise noted.

#### fMRI

Data were preprocessed using AFNI (Cox, 1996). Subject-wise data were transformed to MNI standard space using a two-step linear and nonlinear registration (AFNI’s 3dAllineate and 3dQWarp), and re-sampled to a common rectangular grid at 4 mm resolution using linear interpolation. Data were then spatially smoothed using a Gaussian kernel (full-width-at-half-maximum: 10 mm; with less smoothing IVC were lower and areas above significance more disconnected). After extraction of the brain volume, 32,798 voxels were retained. Volumetric data were further mapped to the cortical surface of the ‘New York Head’ (Huang et al., 2016), a high resolution anatomical template extending the ICBM152 head (Fonov et al., 2011), based on minimum Euclidean distance. This resulted in the selection of 16,037 grey matter voxels (8,080 in the left hemisphere) to be analyzed.

#### ECoG

T1-weighted MR images were acquired from each subject both before and after the implantation of electrodes. Electrodes were localized on the individual cortical surfaces using a combination of manual identification in the T1 images, intraoperative photographs, and a custom MATLAB tool based on the known physical dimensions of the grids and strips (Yang et al., 2012). Subsequently, the individual-subject T1 images were non-linearly registered to an MNI template using the DARTEL algorithm via SPM (Ashburner, 2007), and the same transformation was applied to map individual electrode coordinates into MNI space. Out of 573 artifact-free labeled and MNI-registered electrodes used in Honey et al. (2012), only those 511 located in the left hemisphere were retained (see Fig. S1 for a depiction of the electrode locations). Data of all five subjects were pooled and subsequently treated as one single dataset.

#### EEG

For plotting purposes, electrode coordinates were aligned with the surface of the New York Head (Huang et al., 2016) using MNI coordinates provided by EEGLAB (Delorme et al., 2011) (see Fig. S1 for electrode locations). Source reconstruction (see EEG source modeling) was conducted in order to map EEG activity to cortical anatomy in MNI standard space.

### Data preprocessing

#### fMRI

Functional data were slice-time and motion corrected (AFNI’s 3dvolreg). The remaining data were linearly detrended, and high-pass filtered at 0.01 Hz using the discrete Fourier transform (DFT) and its inverse. In addition to the subject-level data, a grand-average (GA) was computed by averaging across subjects.

#### ECoG

Data were preprocessed using mean regression and down-sampling to 400 Hz (see procedures described in Honey et al., 2012). In order to facilitate comparisons between ECoG and EEG data, ECoG data were further decimated to 256 Hz after appropriate low-pass filtering using anti-aliasing finite impulse response filters (8th order Chebyshev Type I, MATLAB’s resample function).

#### EEG

Data were decimated to 256 Hz using MATLAB’s resample. Data were then high-pass filtered at 0.5 Hz and notch-filtered between 59 and 61 Hz using third order Butterworth infinite response (IIR) filters. Signal from six electrooculogram (EOG) channels was removed from all EEG channels through multivariate linear regression and subtraction, i.e. noise canceling using standard procedures (Parra et al., 2005); code available from Cohen and Parra (2016). Artifact channels, defined as those with unusually small or large standard deviations (SD < 1 μ V or SD > 50 μ V or SD > mean_ch_(SD) + 2 SD_ch_(SD), where SD_ch_ are mean and standard deviations over channels, respectively) were set to zero. This, however, affected only one channel per subject on average and had a negligible impact on the overall results. A grand-average dataset was computed by averaging the preprocessed EEG time courses (here referred to as ‘raw’ as opposed to log band-power time series (see Calculation of EEG/ECoG band-power) across subjects.

In all datasets, the first 15 and the last 13 seconds were excluded, leaving a period of 297 seconds to be analyzed. The initial time is typically removed to allow the T1 saturation to reach a steady state and avoid any potential evoked responses due to scanning onset. In this paradigm we also wanted to remove a strong visual, non-specific response. There was also a brief blank screen period at the end of the recording. The number of EEG/ECoG samples at 256 Hz sampling rate was T = 76,032, while the number of fMRI samples (TR’s) at 0.67 Hz was T = 198.

### EEG source modeling

For source analysis, EEG data at 32 channels were mapped to 2,004 locations covering the entire cortical surface (1,002 in the left hemisphere) by inverting a precise standardized volume conductor model of current flow in an average human head (Huang et al., 2016). This step was performed on grand-average signals for analyses of raw broad-band EEG time courses, while it was performed separately for each subject for analyses of the log-power of brain oscillations (see below). Prior to source imaging, data and head model were transformed into common average electrical reference. The inversion was carried out using eLORETA (Pascual-Marqui, 2007). The regularization parameter λ was adjusted on grand-average raw EEG data using two-fold cross-validation. To this end, electrodes were split into two subsets comprising 19 and 13 electrodes, respectively, both covering the whole scalp. Sources were first estimated based on the subset of 19 electrodes for 11 logarithmically spaced choices of the regularization parameter λ. The resulting source activity was projected back to the scalp using the volume conductor model. The resulting scalp potentials were evaluated at the 13 remaining channels, and the discrepancy to the potentials actually recorded at those electrodes was measured in terms of the mean-squared error (MSE). Using the value of λ that minimized this MSE, sources were re-estimated based on all 32 electrodes. This choice of λ was used in all subsequent source analyses, including band-power analyses in source space.

Three-dimensional source current estimates were further reduced to scalar activations. This was done by projecting the current vector using location-specific 3D projection vectors that were designed to maximize the correlation of the grand-average raw EEG signal between the first and the second viewing (Dmochowski et al., 2012). These projection vectors were computed once and used throughout all source analyses.

### Calculation of EEG/ECoG band-power

In addition to the raw broad-band EEG and ECoG signals (preprocessed as described above), we also calculated the instantaneous amplitude in five common frequency bands. These bands included θ (4–8 Hz), α (8–12 Hz), β (12–28 Hz), γ (28–56 Hz) and a high-frequency band (HF, 64–116 Hz). subject-level data were filtered in each band using third-order Butterworth IIR filters. We applied the Hilbert transform to obtain the complex-valued analytic signal, the absolute value of which provides the instantaneous band-amplitude. For single-subject analysis (see Fig. S2 B, C), the logarithm was applied directly. Grand-average instantaneous log band-power was obtained by taking the mean of the subject-wise squared amplitudes across subjects, and applying the logarithm on the mean. Additional down-sampled datasets were obtained by reducing the EEG and ECoG band data to the fMRI sampling rate of 0.67 Hz using resample.

### Inter-Viewing correlation (IVC) within imaging methods

Raw and log band-power EEG data (at the level of 32 sensors as well as at 2,004 source locations) were linearly detrended. Correlations between time series recorded during the first and second movie viewing were assessed using the Pearson product-moment correlation coefficient, yielding one interviewing correlation (IVC) coefficient per channel and imaging method.

### Mappings between fMRI, ECoG and EEG spaces

In order to study correlations *across* imaging methods we selected channels (electrodes/cortical locations) that were co-located. The procedures to match channels between methods as well as to harmonize the temporal scale of the data for each pair of methods are outlined below.

#### ECoG-fMRI

For each of the 511 ECoG channels, corresponding BOLD activity was defined as the average of the activity of all cortical fMRI voxels within a 6 mm radius around that ECoG electrode. The down-sampled ECoG log band-power data at 0.67 Hz sampling rate were convolved with a canonical hemodynamic response function (HRF, SPM package, Penny et al., 2011). ECoG data were then detrended, high-pass filtered at 0.01 Hz using the Fourier transform, and cropped to the common 297 s interval.

#### ECoG–EEG

EEG channels pick up activity from the entire brain as a result of the spread of neuronal electrical activity in the head. It is, therefore, impossible to associate individual EEG channels with ECoG/fMRI counterparts, which is why we worked on EEG source estimates obtained as described above. Source locations were assigned to the closest ECoG electrode based on minimum Euclidean distance.

#### fMRI–EEG

EEG source log band-power traces at 0.67 Hz sampling rate were convolved with a canonical HRF, detrended, high-pass filtered at 0.01 Hz, and cropped. EEG sources were interpolated to match fMRI voxel locations based on minimum Euclidean distance.

### Inter-method correlation (IMC) analysis

For each of the three pairs of measurement techniques, correlations between the appropriately mapped time courses (averaged across the two viewings) were assessed using Pearson’s product-moment correlation, yielding one inter-method correlation (IMC) value per channel and pair of imaging techniques.

### Statistical significance of correlations

As neurophysiological time series are auto-correlated, correlations between them cannot be assessed using standard analytical tests assuming independent and identically distributed samples. Instead, the distribution of observed correlations under the null hypothesis of zero true correlation needs to be estimated empirically. Correlation values r obtained on original and surrogate data were mapped to the interval [-∞,∞] using the Fisher z-transform ρ = atanh(r), where atanh is the hyperbolic tangent. For each channel, we confirmed that the null distribution of the z-transformed correlation scores is consistent with a Gaussian distribution using the Kolmogorov-Smirnov test (p < 0.05). Means μ_0_ and standard deviations σ_0_ estimated from surrogate data were used to standardize correlation coefficients observed on the original data, yielding z-scores z = (ρ-μ_0_)/σ_0_. We derived p-values assuming that these z-scores are standard normal distributed under the null hypothesis. We demonstrate the exactness of this approach empirically with numerical simulations in Testing for significant correlation between auto-correlated time series: simulation.

As we did not expect any negative inter-viewing correlations, right-tailed tests were used for all IVC analyses. In contrast, inter-method correlations were assessed using two-tailed tests, except for correlations involving ‘raw’ EEG or ECoG signals, the polarity of which depends on recording and inverse source reconstruction parameters, and is essentially arbitrary, as is the sign of the resulting inter-method correlations. Consequently, we tested the absolute value of the IMC in these cases using a one-tailed test. For each analysis, the false discovery rate (FDR) was controlled at level q = 0.05 by applying the Benjamini-Hochberg correction (Benjamini and Hochberg, 1995) across all voxels. Only z-scores that were significant after FDR correction are reported.

### Visualization

Color-coded r-values and z-scores were rendered onto the surface of either the outer head surface or the cortical surface of the ‘New York Head’ (Huang et al., 2016). EEG sensor-space results were projected onto the outer head surface using a spherical harmonics expansion interpolation. fMRI results as well as results of EEG–fMRI IMC analyses were mapped to the cortical surface using to the nearest-neighbor rule, where distance was measured along the geodesics of the cortical manifold. ECoG results as well as results of ECoG–fMRI and EEG–ECoG IMC analyses were mapped onto the cortical surface by coloring all surface nodes within 6 mm distance of each ECoG channel. Cortical surfaces were smoothed for display purposes. Electrodes/voxels with z-scores significant after FDR correction are shown in color, while non-significant locations are shown in gray. Identical r-values/z-scores are displayed in identical colors in all plots throughout the paper except for Fig. 1 D and for Fig. S2. To cover the entire range of values observed, we used nonlinear saturating color scales. For each analysis, a colorbar is plotted that shows the range of attained r-scores/significant z-scores.

## ACKNOWLEDGEMENTS

SH was supported by a Marie Curie individual International Outgoing Fellowship (grant No. PIOF-GA-2013-625991) within the 7th European Community Framework Programme. PD, LCP, and SH were supported by Defense Advanced Research Projects Agency (DARPA) contracts W31P4Q-13-C-0038 and W911NF-14-1-0157. UH, CJH and LCP acknowledge support from the National Institutes of Health (NIH, 1DP1HD091948-01 award to UH, MH111439-01 subaward to CJH, R01-NS095123 and R01MH111439 awards to LCP). CJH further gratefully acknowledges a Research Fellowship awarded by the Sloan Foundation. We would like to thank Ricardo Henriques for providing this 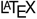 template.

## APPENDIX Testing for significant correlation between auto-correlated time series: simulation

We assessed the validity of the statistical test for non-zero correlations between auto-correlated time series introduced in the Statistical significance of correlations section. Two independent time series were generated either as univariate third-order linear auto-regressive (AR) processes, as Gaussian-distributed pink noise, or as squared (non-Gaussian distributed) pink noise. The number of samples was set to either N = 198 or to N = 76,032, amounting to a recording of 297 s length at the fMRI and EEG/ECoG sampling frequencies, respectively. For each pair of time series, 100 surrogate datasets were constructed using phase randomization, and a p-value was derived. An alternative p-value was derived using MATLAB’s corr() function under the assumption that samples are independent. For this approach, correlation scores were transformed into a Student-t distributed test statistic, which gave rise to an analytic solution for the p-value.

Each experiment was repeated 2,000 times. From the distribution of the p-values we derived the empirical false-positive rate (FPR) as a function of the required FPR (alpha-level). For a statistical test to be exact, it is important that the relation between the two is close to identity, while undershoots of the empirical FPR are generally more tolerable than inflated FPR’s. Results shown below (Fig. A1) indicate that the statistical test based on surrogate data is faithful to the desired alpha level for all possible FPR’s and in all tested scenarios. In contrast, the standard analytic test is characterized by highly inflated FPR’s in the alpha ranges of interest in practice (α = 0.01, α = 0.05). This behavior is more pronounced for pink noise than for auto-regressive processes, and for longer compared to shorter time series. In extreme cases, FPR’s of up to 92% (pink noise data, N = 76,032) are observed at a nominal alpha level of 5%.

**Fig. A1.**
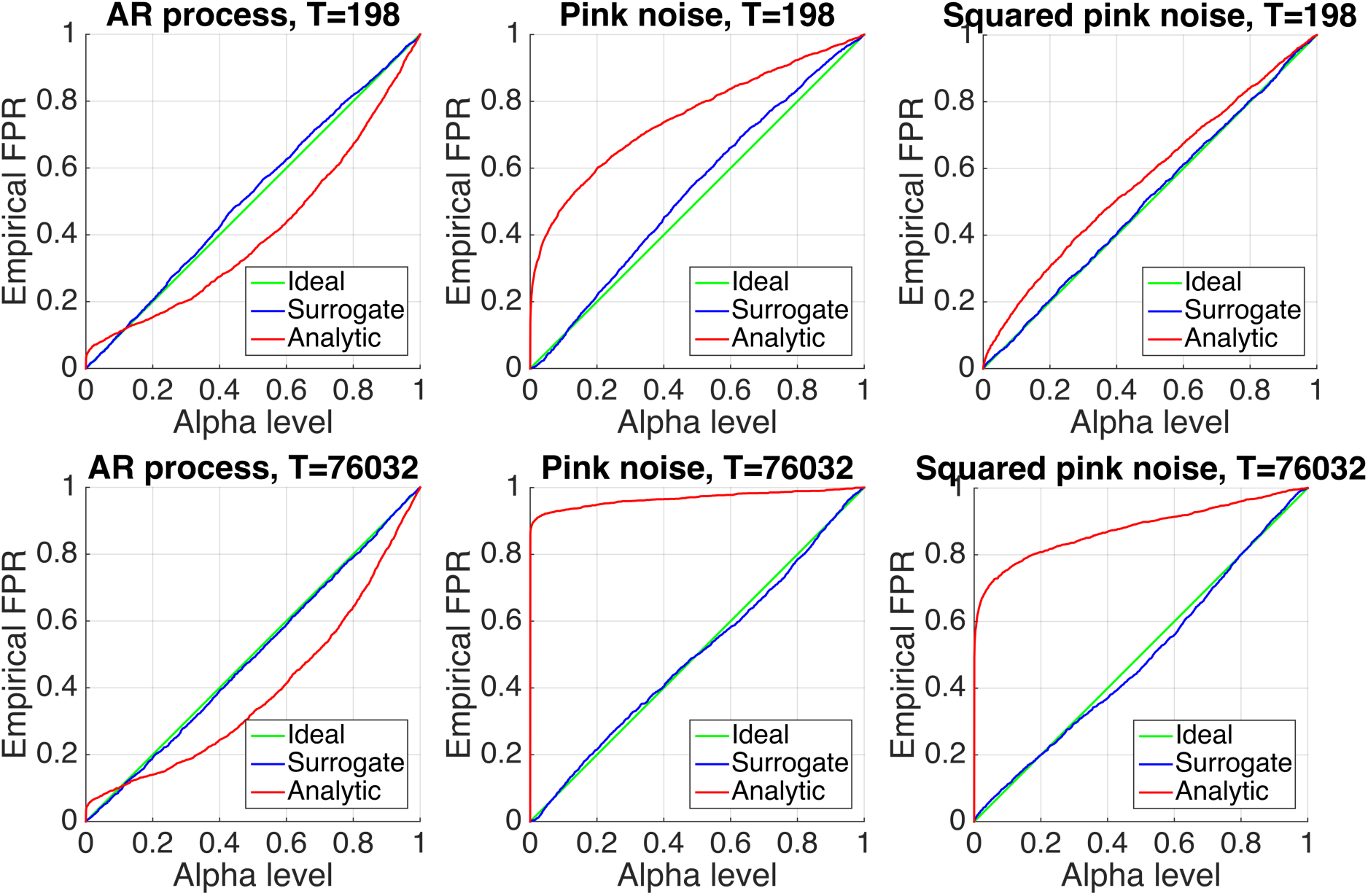
**Comparison of two approaches to test the statistical significance of correlations** between auto-correlated time series: the surrogate data based approach used throughout this paper (see Statistical significance of correlations section) and the conventional analytic approach assuming independent samples. The surrogate based approach leads to empirical false-positive rates (FPR) that are close to the desired FPR (alpha level) regardless of the sample size or temporal dynamics of the time series, while the conventional approach leads to strongly inflated FPR’s in the practically relevant alpha ranges (α = 0.01, α = 0.05) in all cases.

## SUPPLEMENTARY FIGURES

**Fig. S1.**
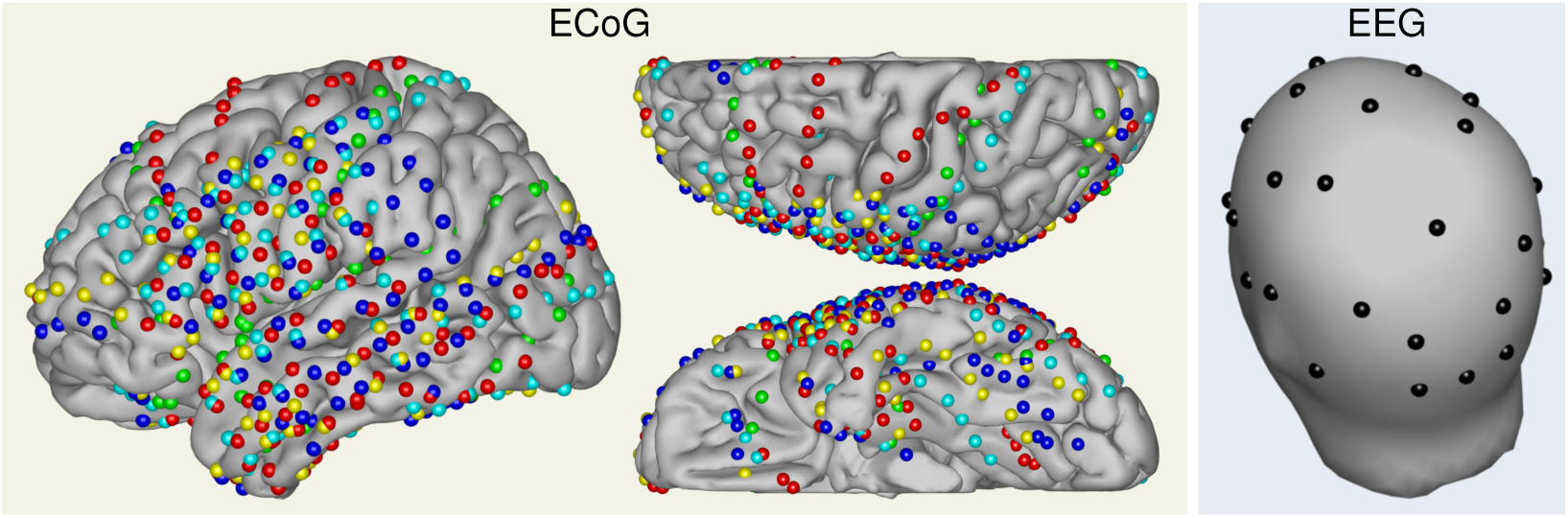
Location of ECoG and EEG electrodes on the scalp/cortical surfaces. ECoG electrodes are drawn in a different color for each of the five subjects.

**Fig. S2.**
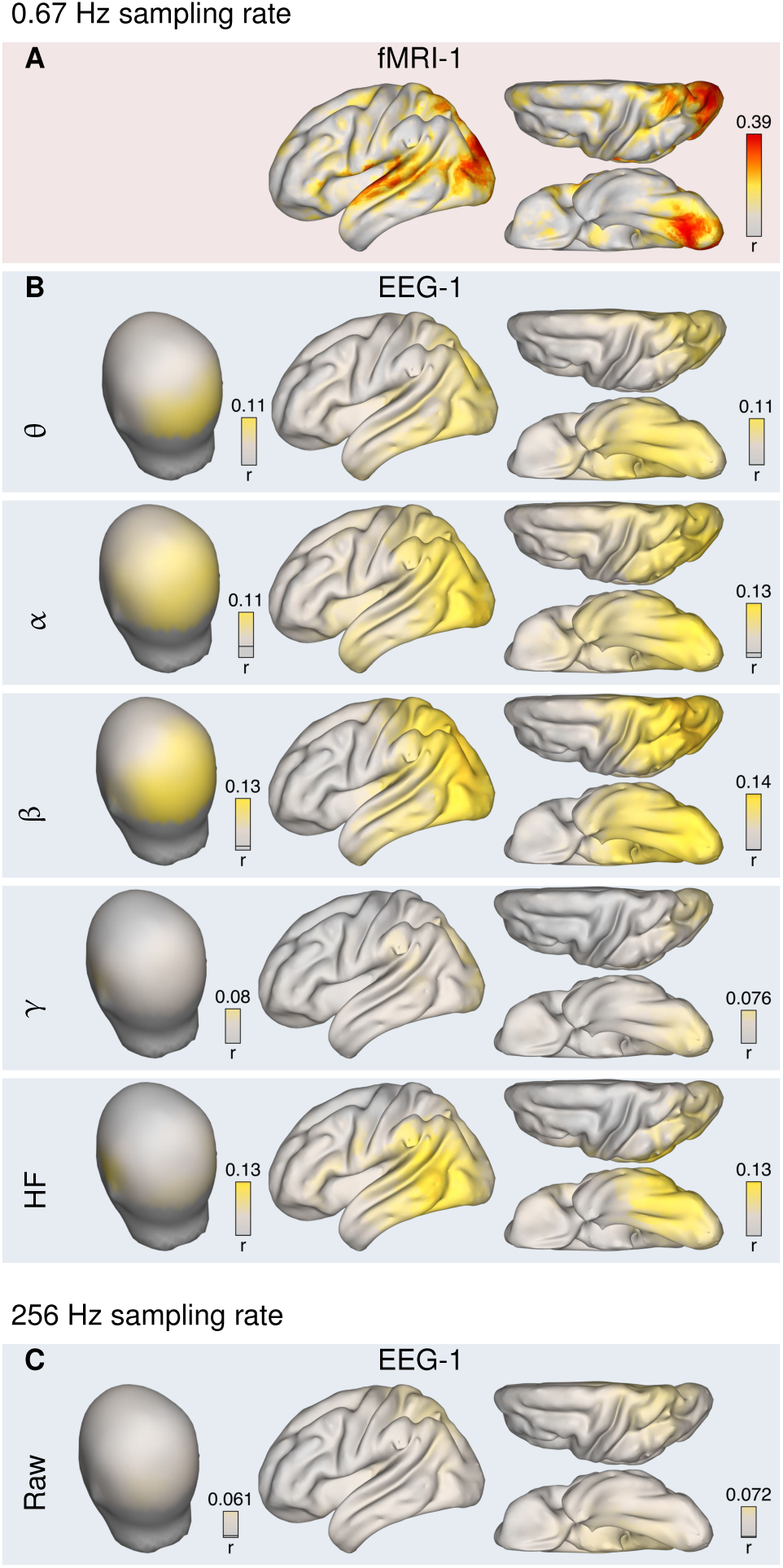
Inter-viewing correlations observed within single subjects for EEG and fMRI. IVC scores, *r*, were computed in each subject, and averaged across subjects. No statistical testing/thresholding was performed. Observed IVC scores are smaller than those observed on grand-averaged data (c.f., Fig. 1, but notice the different color scale here), yet exhibit similar topographies.

**Fig. S3.**
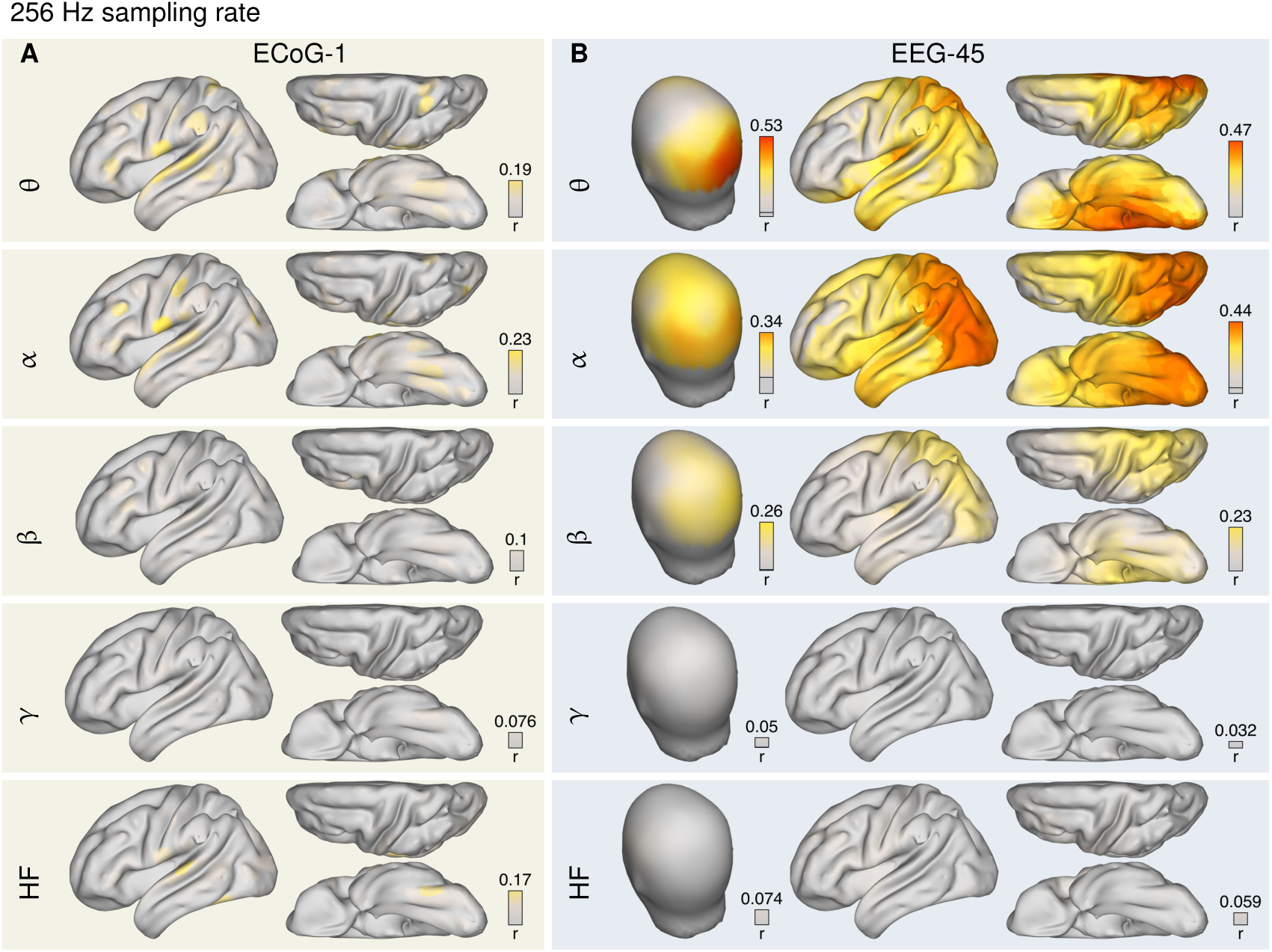
**Inter-viewing correlation** of EEG/ECoG band-power computed **at 256 Hz sampling rate**, i.e., without low-pass filtering the instantaneous log band-power. Observed IVC values are smaller than for down-sampled data, but exhibit similar spatial topographies.

**Fig. S4.**
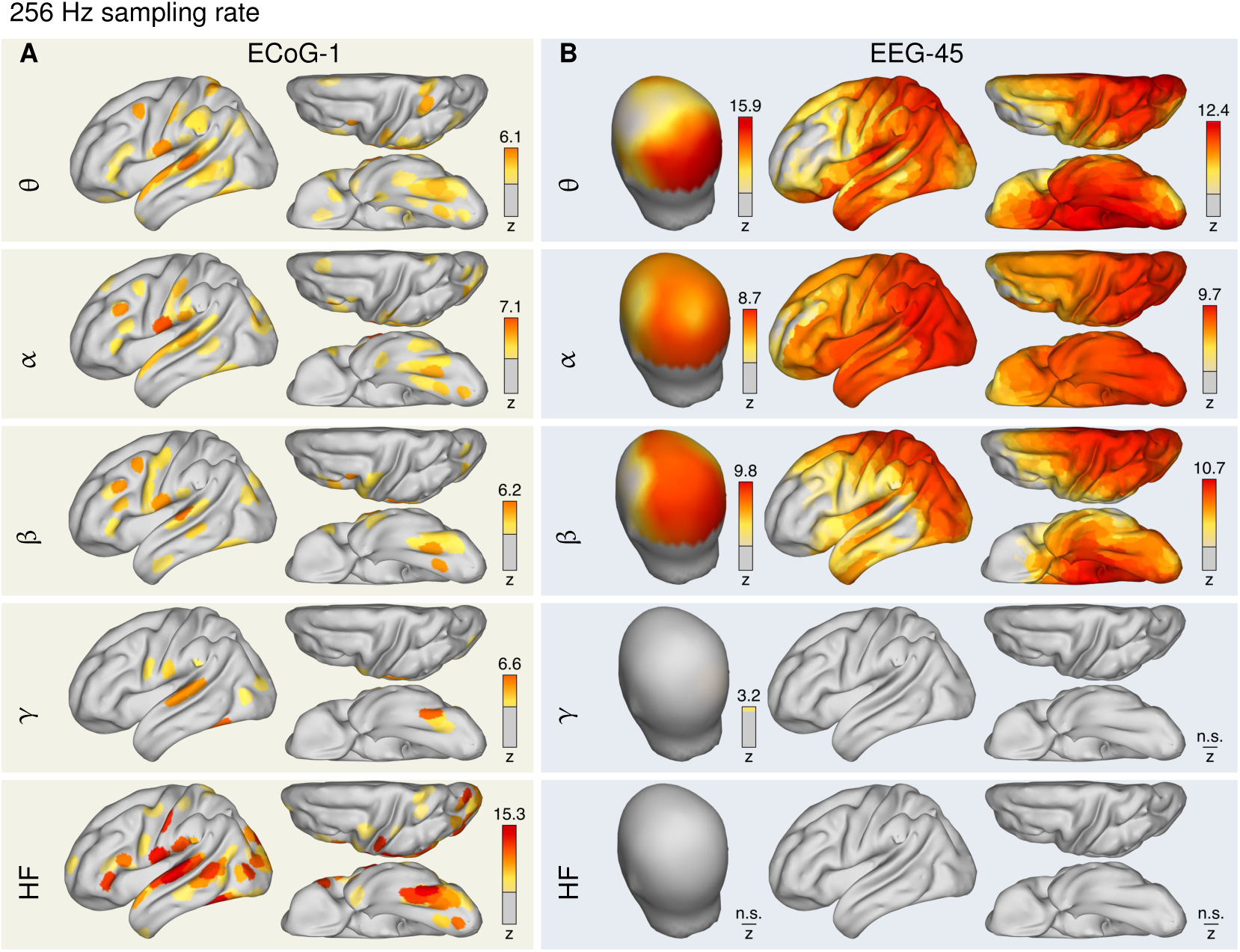
**Statistical significance** of EEG/ECoG band-power inter-method correlation at **256 Hz sampling rate**. Note that even small IVC values (as, e.g., observed for high-frequency ECoG data, c.f. Fig. S3) are highly significant.

**Fig. S5.**
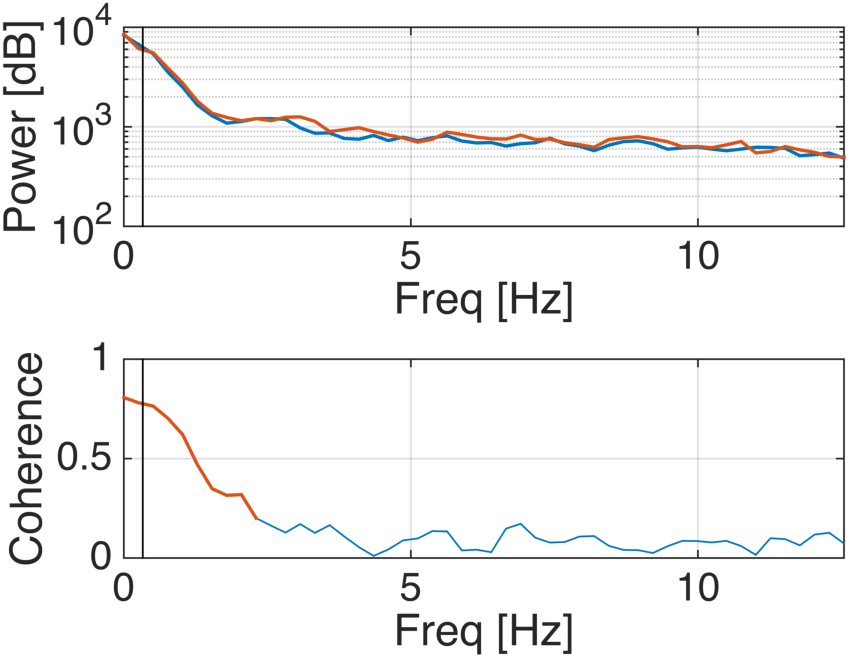
Analysis of inter-viewing coherence (frequency-resolved correlation) of high-frequency (56–116 Hz) ECoG band-power fluctuations for the ECoG electrode exhibiting highest IVC in the HF band. Top panel shows the power spectrum for first and second viewing. The HF power signal is dominated by activity below 2 Hz. The coherence spectrum shows significant correlation (red bold line) also below 2 Hz. The black vertical line indicates the low-pass filter at 0.33 Hz that is applied when sampling at the fMRI resolution of 0.67 Hz. For this signal, an inter-viewing correlation of r = 0.17 (z = 15.3) was observed at 256 Hz sampling rate. After low-pass filtering and down-sampled to 0.67 Hz, IV correlation increased to r = 0.77, but was slightly less significant (z = 10.5). This drop in significance is explained by the fact that the low-pass filter removes correlated signal components that do not exist in random surrogate data. On the other hand, restricting the signal content to the strongest and most correlated portion through the low-pass filter increases inter-viewing correlation, as less correlated signal parts are removed.

**Fig. S6.**
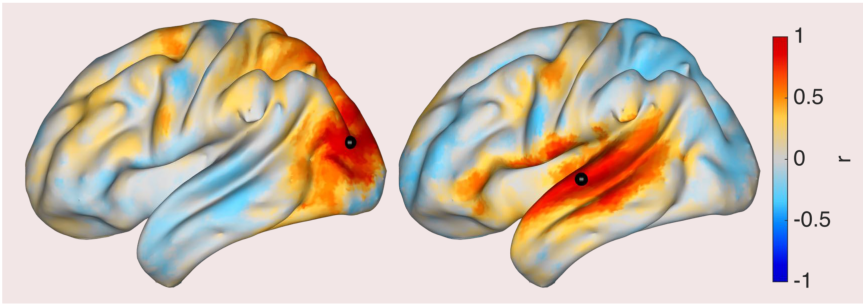
Correlation between different cortical areas within fMRI. Seeds for computing correlations with other fMRI voxel are in occipital (left) and temporal cortex (right), indicated as black dots.

**Fig. S7.**
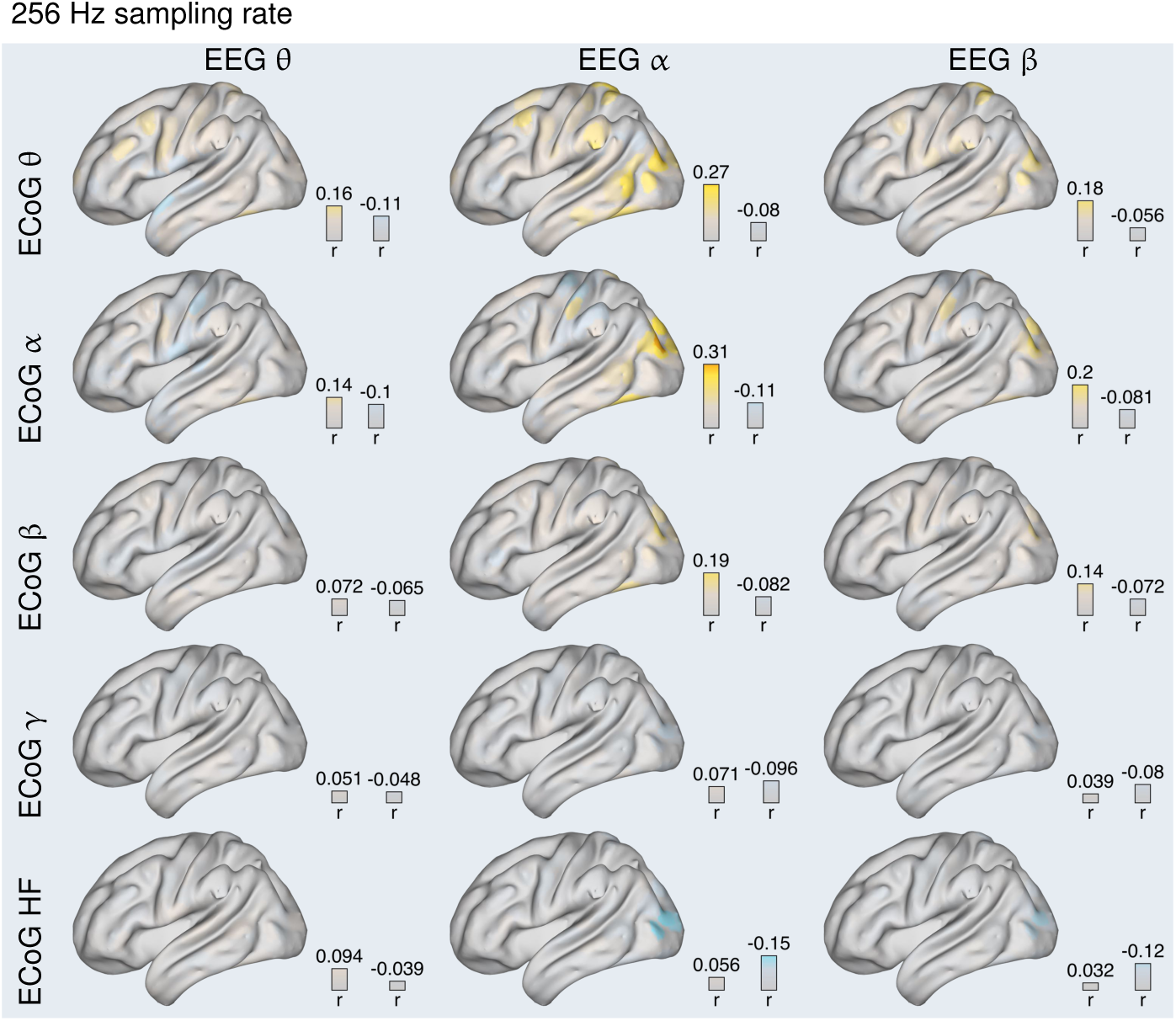
**Inter-method correlation (IMC)** between the instantaneous log-power of EEG and ECoG oscillations in the θ (4–8 Hz), α (8–12 Hz), β (12–28 Hz), γ (28–56 Hz) and high-frequency (HF, 64–116 Hz) bands **at 256 Hz sampling rate**.

**Fig. S8.**
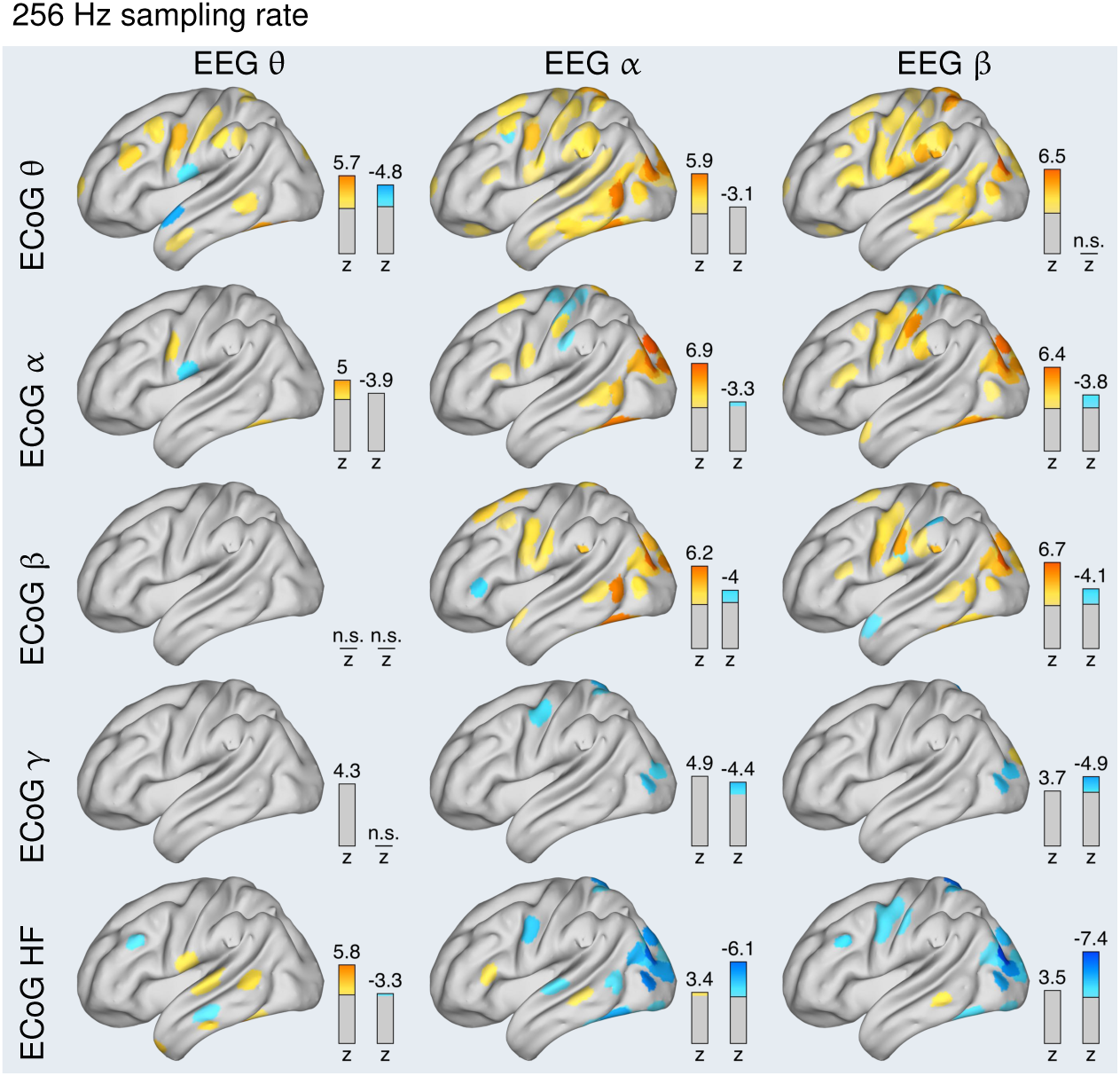
**Statistical significance of inter-method correlations** between EEG and ECoG band-power **at 256 Hz sampling rate**. While correlation values are much smaller than for 0.67 Hz sampling rate, they remain equally significant as compared to null distributions.

